# Cellular, molecular, and therapeutic characterization of pilocarpine-induced temporal lobe epilepsy

**DOI:** 10.1101/2021.01.08.425892

**Authors:** Nicholas D. Henkel, Marissa A. Smail, Xiaojun Wu, Heather A. Enright, Nicholas Fischer, Hunter M. Eby, Paul R. Carney, Robert E. McCullumsmith, Rammohan Shukla

## Abstract

We probed a transcriptomic dataset of pilocarpine-induced TLE using various ontological, machine-learning, and systems-biology approaches. We showed that, underneath the complex and penetrant changes, moderate-to-subtle upregulated homeostatic and downregulated synaptic changes associated with the dentate gyrus and hippocampal subfields could not only predict TLE but various other forms of epilepsy. At the cellular level, pyramidal neurons and interneurons showed disparate changes, whereas the proportion of non-neuronal cells increased steadily. A probabilistic Bayesian network demonstrated an aberrant and oscillating physiological interaction between oligodendrocytes and interneurons in driving seizures. Validating the Bayesian inference, we showed that the cell types driving the seizures were associated with known antiepileptic and epileptic drugs. These findings provide predictive biomarkers of epilepsy, insights into the cellular connections and causal changes associated with TLE and a drug discovery method focusing on these events.

## INTRODUCTION

Epilepsy is one of the most prevalent neurological disorders and is a common end point of several brain pathologies (Stafstrom and Carmant, 2015). Diagnosed as the presence of spontaneous and recurrent seizures, epilepsy is a network-level phenomenon involving changes in neuronal activity, content, and density, leading to re-entrant activation of neurons embedded within brain circuits (DeFelipe, 1999; Jiruska et al., 2010; Pitkänen and Sutula, 2002). Although it is usually acquired following injury to a previously normal brain, familial forms of epilepsy are also known (Berkovic et al., 2006; Christensen et al., 2009).

Temporal lobe epilepsy (TLE) is the most common form of epilepsy, accounting for >40% of adult cases (Blair, 2012; Salzmann and Malafosse, 2012). Animal models that mimic TLE are presently available and, since their establishment, significant progress in the understanding of epilepsy has been achieved (Lévesque et al., 2016). The knowledge gained through these models is helpful in understanding the highly dynamic processes of TLE, such as hyperexcitability and pharmacoresistance (Wang et al., 2003), mostly through electrophysiological techniques, and has helped reduce the number of seizures, but cannot completely cure them (Löscher, 2011). Molecular studies complementing the electrophysiological findings have revealed new molecular targets, identified antiepileptic drug (AED) candidates (Dixit et al., 2016), and confirmed experimental findings implicating the GABAergic system and ion channels in epilepsy (Pfisterer et al., 2020). However, the use of this information in clinical practice is impeded by the lack of studies tracing the cellular origin of these molecular changes and cell-specific drug effects in TLE (Guerrini et al., 2003). Furthermore, existing molecular studies lack discrimination between the upstream (causal) and downstream (consequential) changes associated with epileptogenesis as well as information on their interactions and cellular correlates.

Here, we hypothesized that (1) an integrated analysis of transcriptomics data from bulk and single-cell RNAseq experiments will aid in tracing the cellular origins of TLE, (2) a Bayesian inference-based network biology approach applied to transcriptomics profiles will help identify distinct causal and consequential changes associated with TLE, and (3) drug-based transcriptomics signatures serve as a proxy for the lack of interventional data required to support causal inferences. We predicted key nodes involved in TLE and the cellular correlates of several epilepsy-inducing and antiepileptic drugs.

## RESULTS

### Gene expression changes in the epileptic phenotype

To analyze changes in gene expression associated with pilocarpine-induced TLE, we used RNAseq data generated from 108 epileptic and 103 control adult male Crl:NMRI (Han) FR mice (Srivastava et al., 2018). Principal component analysis clearly segregated the control and epileptic phenotypes (Figure 1A). Indicative of the penetrant nature of the epileptic phenotype, ~75% (11,580/15,300) of the genes in our analysis were differentially expressed in the epileptic vs. the control condition (Up: 5,722 genes; Down: 5,858 genes; FDR < 0.05; Table S1). Biologically significant changes can occur at any level of fold change (St. Laurent et al., 2013), and the number of samples in the present dataset provided sufficient power to detect subtle yet significant (*q* < 0.05) changes. Thus, to determine the abundance of genes at different fold changes, we binned and plotted the DEGs at different thresholds of log-fold change (Figure 1B). The histogram suggests that subtle changes with an absolute log-fold change of <0.2 were most abundant for both up- and downregulated changes; however, they were more prominent in the downregulated direction (Down: 4,156; Up: 2,830). Conversely, distinct changes with an absolute log-fold change of >1 were few among both up- and downregulated changes but were more prominent in the upregulated direction (Up: 269; Down: 20).

**Figure 1:**
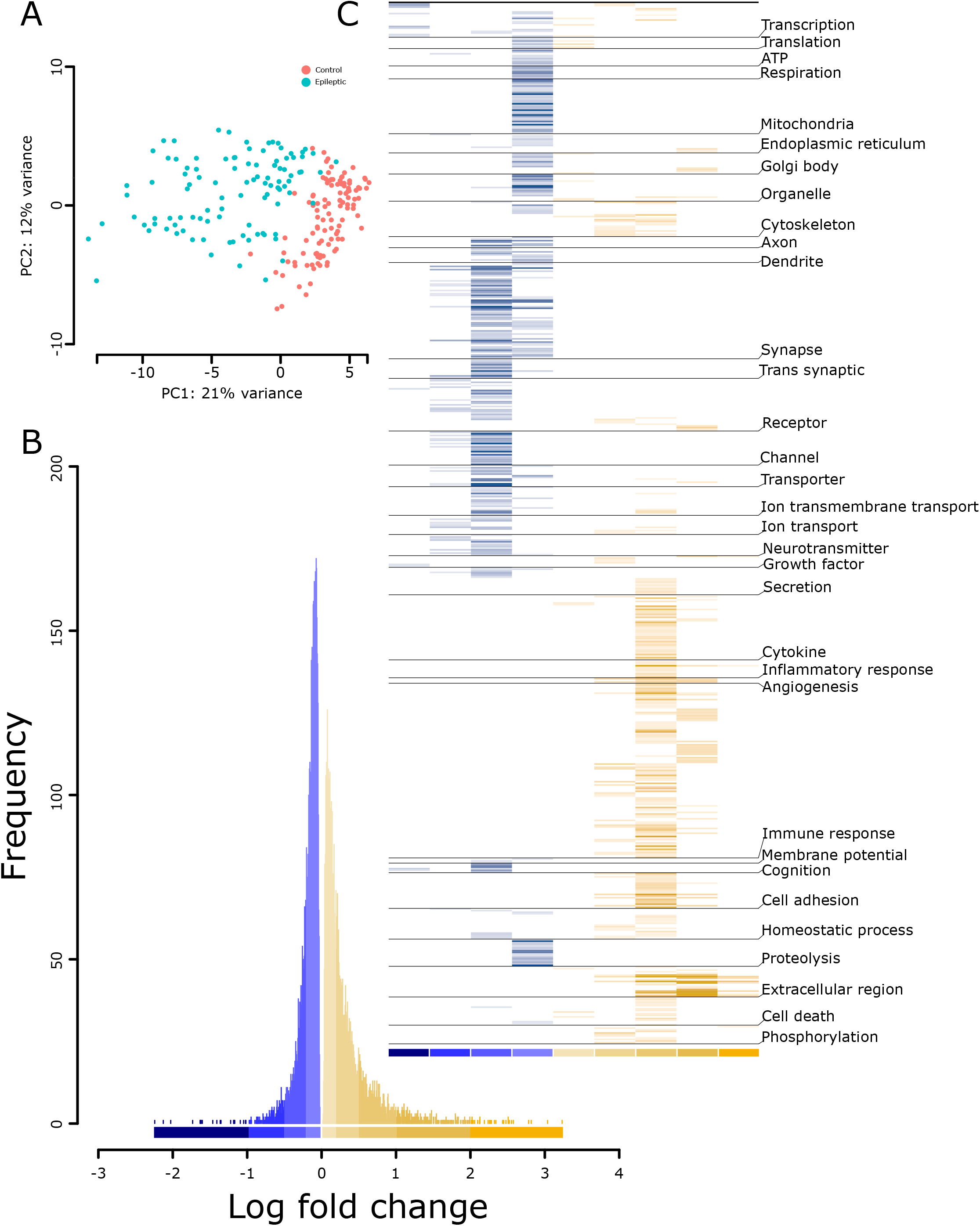
Distinct transcriptomics and pathway profiles of pilocarpine induced TLE in mice. (A) Principal components analysis segregating the transcriptomics profile of the control and pilocarpine induced TLE mice. (B) Frequency of differentially expressed genes binned based on log-fold changes across down-(blue) and up-(yellow) regulated genes. See Table S1 for genes under each bin. (C) Pathway profiles of up- and downregulated genes by log-fold change bin (from B) using Gene Ontology. The -log_10_(q-value) of each pathway was used to plot the heatmap. The lighter-to-darker shades of blue and yellow indicate increasing significance of down- and upregulated pathways, respectively. See Table S2 for the pathways grouped under each theme.

### Down- and upregulated changes in the epileptic phenotype are biased toward specific biological effects

We explored the biological pathways associated with different bins of DEGs. All identified pathways (*q* < 0.05) were clustered based on functional themes (Figure 1C, Table S2).

#### Downregulated pathways

Highly downregulated genes (log-fold change <−1) corresponded to a few pathways and were associated with themes related to *transcription, membrane potential, receptors, and growth factors*. Moreover, 8/13 pathways in this category predominantly featured *transcription factor binding*, suggesting an overall decrease in regulatory events. Intermediately downregulated genes (log-fold change ranging between −1 and −0.5) were largely related to neuronal signaling. Furthermore, 45 pathways in this bin were associated with *synapse, receptors, channels, ion transport, neurotransmitter*, and *G-protein-coupled receptors*. Subtly downregulated changes (log-fold change between −0.5 and −0.2) had 180 pathways and, similar to the prior bin, were predominantly associated with *synapse*- and *signaling*-related themes. The most subtle changes (log-fold change between 0 and −0.2) had 174 pathways associated with *ATP production, mitochondria, proteolysis, transcription*, and *translation*, indicating impairment in energy balance and basic cell functionality in TLE. Interestingly, signaling-associated themes (*axon, dendrite, synapse, receptors, ion channels, ion transport, neurotransmitter*, and *membrane potential*) were dominant in all the downregulated pathway bins, suggesting the altered and widespread impact of neuronal activity in TLE.

#### Upregulated pathways

Highly upregulated genes (log-fold change >2) corresponding to 10 pathways were almost exclusively related to *extracellular matrix*. Other pathways in this bin were associated with *phosphorylation, growth factors*, and *inflammation*. Genes with a log-fold change of 1– 2 corresponded to 68 pathways largely related to *immune system*, while also featuring *extracellular matrix, angiogenesis*, and *organelle activity* (including of Golgi bodies and the endoplasmic reticulum). Intermediately upregulated genes (log-fold change, 0.5–1) corresponding to 232 pathways were largely enriched in *immune system* processes and *extracellular matrix*. We also identified *apoptosis-*, *homeostasis-*, *cell adhesion-*, and *cytoskeleton*-associated pathways. Subtly upregulated changes (log-fold change, 0.2–0.5) shared most themes with the prior bin (0.5–1). There were very few most subtly upregulated pathways (log-fold change, 0–0.2; n + 17), which were related to *transcription, translation, organelles, cytokine production*, and *apoptosis*.

Overall, the down- and upregulated changes in the epileptic phenotype were biased toward specific biological effects. The downregulated pathways were mostly associated with intracellular and cell-to-cell *signaling*, while the upregulated pathways were mostly associated with the extracellular matrix, immune response, and homeostatic changes.

### TLE is associated with subtle molecular changes

Not all DEGs contributed to or informed about the epileptic phenotype. To identify epilepsy-predicting genes, we deployed a sparse classifier that used a minimal set of variables (genes) to predict TLE. Using 119 genes (Table S3; Up: 59 and Down: 60; termed “predictive gene set” hereafter), the classifier could distinguish the withheld control and epilepsy samples with 98% accuracy. For validation, we looked for their enrichment in gene sets associated with various disease phenotypes using the Enrichr database (Kuleshov et al., 2016). The strongest signal stemmed from *genetic reflex epilepsy* (*q* = 6.53 × 10^−37^), *epileptic encephalopathy* (*q* = 1.5 × 10^−35^), and *familial TLE* (*q* = 4.4 × 10^−8^).

Furthermore, we looked for predictive gene enrichment in 1) different DEG bins (Figure 1B), 2) GO terms, and 3) hippocampus cell types/subfields (Figure 2A). Within different DEG bins, the predictive genes were enriched in up- and downregulated bins of subtle (Down: lfc + −0.5 to −0.2, P < 0.3 × 10^−32^; and Up: lfc < 0.2, P < 0.0017; lfc + 0.2 to 0.5, P < 0.0013) to moderate (lfc + −1 to −0.5, P < 0.90 × 10^−3^) log-fold changes. For GO terms (Table S3), similar to the pathway analysis of DEG bins (Figure 1C), the downregulated predictive genes were enriched in pathways associated with cell-to-cell *signaling* involving both *presynapse* (*q* < 1.86 × 10^−18^) and *postsynapse* (*q* < 2.20 × 10^−21^), whereas the upregulated predictive genes were enriched in pathways involving *neurogenesis* (*q* < 3.03 × 10^−8^), *regulation of gliogenesis* (*q* < 5.23 × 10^−3^), and *cellular homeostasis* (*q* < 5.79 × 10^−4^). Finally, within the hippocampal cell types/subfields (Figure 2, details below), the downregulated predictive genes were enriched in cells of the dentate gyrus (cluster-0, q + 6.8 × 10^−09^) and pyramidal neurons of the Cornu ammonis 1 (CA1)|CA2 (cluster-03, q + 3.60 × 10^−28^) and CA3 (cluster-04, q + 6.94 × 10^−07^) subfields, whereas upregulated predictive genes were enriched in the dentate gyrus (cluster-0, q + 7.28 × 10^−07^), astrocyte (cluster-02, q + 4.31 × 10^−26^), and pyramidal neurons of CA1|CA2 (cluster-03, q + 6.3 × 10^−3^) subfields.

**Figure 2:**
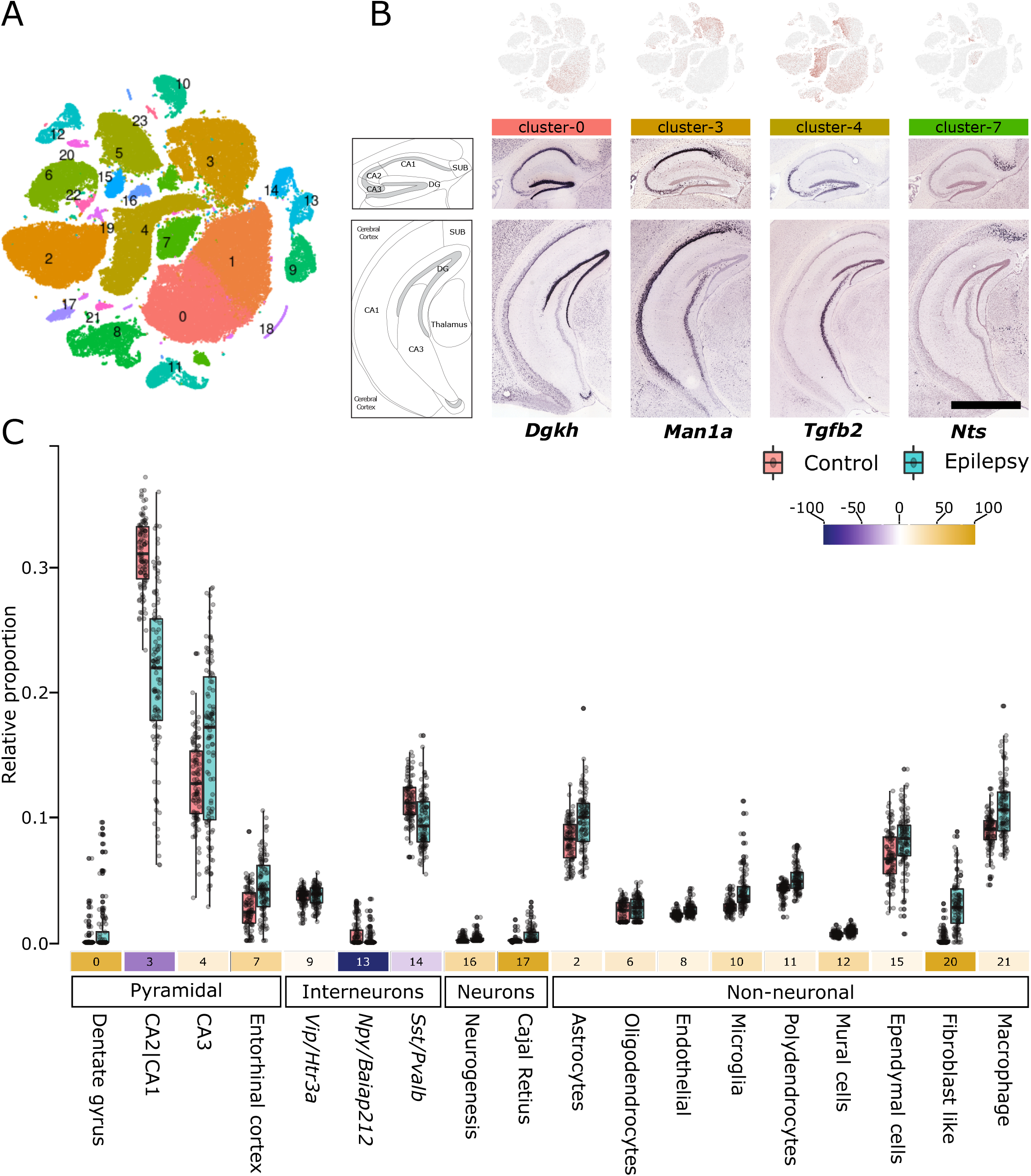
Cell-type deconvolution of TLE RNAseq data reveals altered neuronal and non-neuronal cell proportions. (A) Clustering of mouse hippocampus single-cell RNAseq data from Gene Expression Omnibus (GSE116470). See Figure S1 for cluster characterization and Table S4 for cluster-specific genes. (B) Cell-type clusters representing pyramidal neurons were enriched in different hippocampal regions and subfields. In situ hybridization data (Allen brain atlas) of genes (discrete markers, Table S4) representative of the pyramidal-neuron-specific cluster-0 (*Dgkh*), cluster-3 (*Man1a*), cluster-4 (*Tgfb2*), and cluster-7 (*Nts*) are shown. Scale bar, 1678 micron. (C) Proportional differences of cell-type clusters in the control and epileptic cohorts. The Y-axis shows the relative proportion of different cell-type clusters. The number below each box plot represents the cluster number from (A), and the colors in shades of blue and yellow indicate a decrease or increase in the percentage of relative cell proportions, respectively. All changes in proportions were significant (P < 0.001; Wilcoxon test). See Table S4 for additional details.

Overall, the narrowing down of the highly informative differentially expressed genes using a sparse classifier suggested that subtle-to-moderate fold changes in the expression of a few genes enriched in the astrocytes and pyramidal cell types of the dentate gyrus and hippocampal subfields, can predict various forms of epilepsy.

### TLE is associated with altered neuronal and non-neuronal cell proportions

Although the present dataset (derived from a tissue homogenate) could assess the biological changes associated with TLE, the variability in relative cell proportions and associated biological signals was masked. Single-cell transcriptomics datasets and statistical deconvolution methods can be used to address these issues (Baron et al., 2016; Frishberg et al., 2019; Newman et al., 2019). Using the single-cell transcriptomics dataset for the hippocampus (Saunders et al., 2018), we first generated 24 cell-type-specific clusters (Figure 2A), which were characterized in two stages: (1) globally, based on the presence of known markers associated with different cell types (Figure S1), and (2) locally, based on the top upregulated DEGs associated with the respective cluster. For instance, *Vip/Htr3a-, Npy/Baiap212-*, and *Sst/Pvalb*-positive interneurons represented the top DEGs in clusters 9, 13, and 14, respectively (Figure S1). For pyramidal neurons, the clusters also corresponded to recognizable areas of the hippocampus. For instance, cluster-0, cluster-3, cluster-4, and cluster-7 belonged to the dentate gyrus, CA2|CA1 subfields, CA3 subfield, and entorhinal cortex of the hippocampus, respectively (Figure 2B). Using a support-vector-machine-based method (Aaron M. Newman et al., 2015), highly discriminative markers for each cluster (Figure S2, Table S4) were used to deconvolve their relative proportions in the control and epileptic bulk RNAseq samples. Significant differences in proportion were detected for all clusters (Figure 2C), and interesting patterns of changes were noted. First, all non-neuronal cells showed an increased relative proportion in epilepsy vs. control samples. Second, neuronal cells showed both an increase and a decrease in relative proportion. In particular, pyramidal neurons, with the exception of those belonging to the CA2|CA3 subfield, showed an increase in relative proportion, and interneurons, with the exception of *Vip/Htr3a*-positive ones, showed a decrease in relative proportion. Third, pyramidal neurons of the CA1|2|3 subfields showed the highest variation among all cell types.

Cell-specific enrichment, similar to deconvolution (Figure 2C), requires cell-specific marker genes and can be used as a proxy to deduce cell-type proportion at lower resolutions (Shen-Orr and Gaujoux, 2013). To independently confirm our deconvolution analysis, we performed cell-type enrichment within each DEG bin using reported hippocampal cell-specific markers (Mancarci et al., 2017) (Figure 3A). Confirming the changed proportions, we observed the enrichment of interneurons and non-neuronal cells in down- and upregulated bins, respectively.

**Figure 3:**
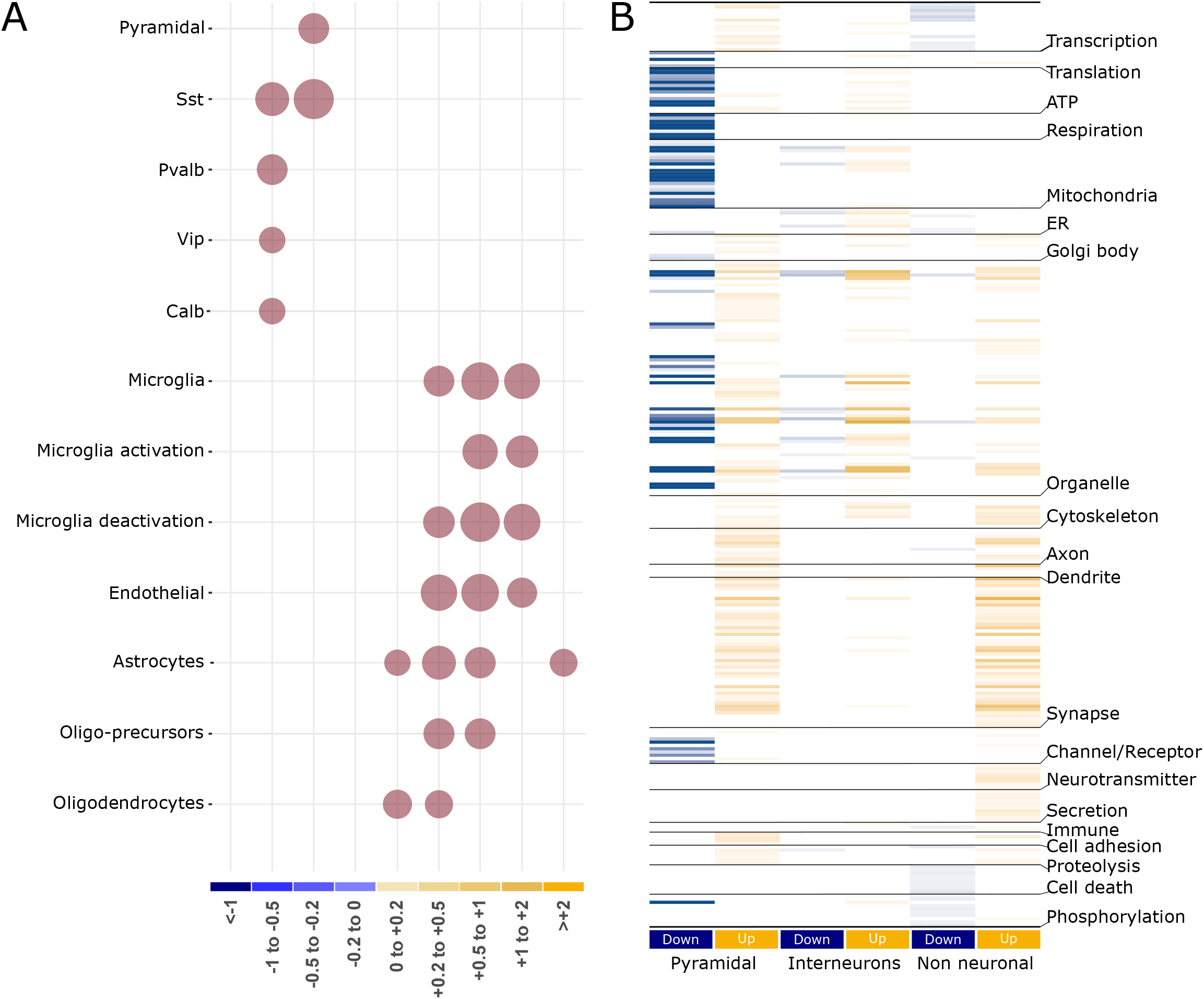
Validation of the cell-type deconvolution (A) and segregation of pathways affected by different cell classes (B). A) Enrichment of cell-specific markers from an independent study within the log-fold change bins of differentially expressed genes. Note that enrichment does not provide detailed segregation of cell type, as observed in Figure 2C. However, reflecting the increased proportion of non-neuronal cells, their enrichment is observed in the bins associated with upregulated genes. The size of the circles is proportional to the -log_10_ value of the enrichment q-value. Significant results (q-value < 0.05) are plotted exclusively. B) Pathways and themes (same as Figure 1C) affected by cell-type clusters representing pyramidal, interneuron, and non-neuronal cells. The log_10_(q-value) of each pathway was used to plot the heatmap. The lighter-to-darker shades of blue and yellow indicate increasing significance of down- and upregulated pathways, respectively. See Table S5 for additional details.

To identify genes that are affected by the altered pyramidal neurons, interneurons, and non-neuronal cells, we regressed out the variability associated with these cell proportions from the differential expression analysis. We found that 1,749 (Up: 709, and Down: 1,040), 2,352 (Up: 1,141, and Down: 1,211), and 1,510 (Up: 761, and Down: 749) genes showed a further reduced *P*-value after regressing the pyramidal neurons, interneurons, and non-neuronal cell proportions, respectively (Table S5). Mapping the themes observed in the prior analysis (Figure 1C) to the pathways associated with these genes (Figure 3B and Table S5) showed that pyramidal neurons prominently influenced the downregulation of bioenergetics (*ATP, respiration, and mitochondria*)-, organelle-, and receptor/channel-associated pathways and the upregulation of synaptic functions; interneurons affected at a low level the downregulation of organelle (*mitochondria and endoplasmic reticulum*)- and the upregulation of cytoskeleton-related pathways, while non-neuronal cells affected the downregulation of transcription-, cell-death-, and phosphorylation-, and the upregulation of cytoskeleton-, axon-, dendrite-, and neurotransmission-related pathways (Figure 3B).

Together, deconvolving the cellular origin of gene expression consistent with the penetrant nature of TLE revealed alterations in all cell types. Opposing changes were observed in neuronal and non-neuronal cells. All non-neuronal cells exhibited an increased proportion, interneurons (with the exception of those expressing *Vip/Htr3a*) showed a decreased proportion, and pyramidal neurons (particularly in CA subfields) showed the highest variation in proportion.

### Investigating causality using a Bayesian gene network analysis

Although the pathway and deconvolution analyses provided biological and cellular insights, they did not inform about events occurring up- and downstream of the disease state. Therefore, we summarized the expression profiles of control and epileptic samples into consensus modules of co-expressed genes using consensus WGCNA (Langfelder and Horvath, 2007) and then used the module eigengene to construct a Bayesian network modeling the probabilistic dependencies between modules as directed acyclic graphs (Zhang et al., 2013) (DAGs; Figure 4A, B; the modules in DAG are termed “nodes” hereafter). To infer the TLE-associated up- and downstream events, we anchored a binary variable associated with sample labels (i.e., control and epileptic) to the DAG. By design, this variable served as 1) a source, with no parent node above it and the child nodes serving as downstream events, or 2) a sink, with no child nodes below it and the parent nodes serving as upstream events.

**Figure 4:**
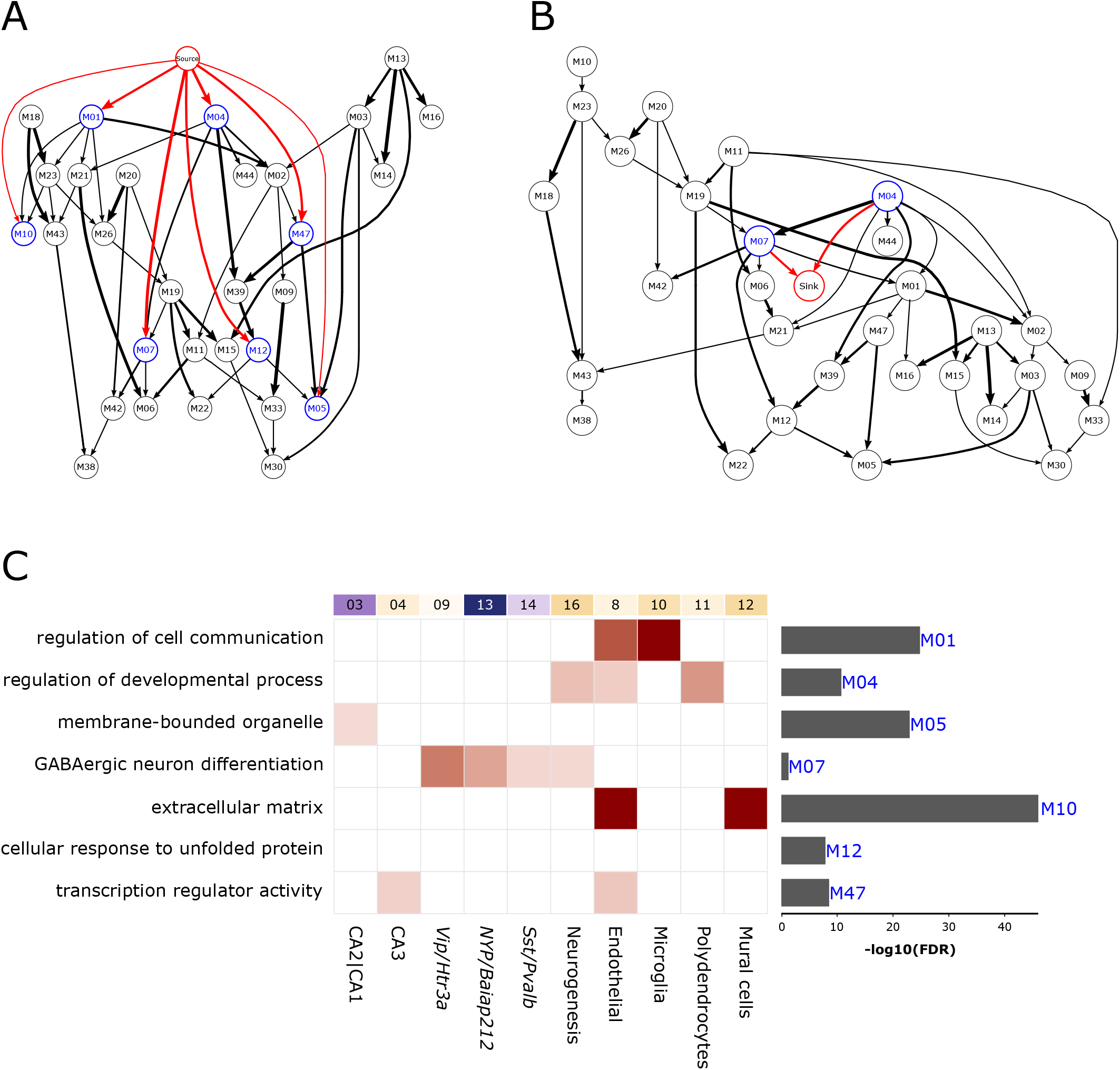
Prioritizing the TLE-associated upstream (putative causal) and downstream (putative consequential) gene modules using a Bayesian network. (A, B) Directed acyclic graph obtained after fitting the Bayesian network to the eigengenes of consensus WGCNA modules, shown as nodes. The edges represent probabilistic dependencies between the nodes from early to later (referred as parent-to-child) associations. Source and sink nodes constitute a categorical variable represented by sample labels (i.e., control and epileptic). Source node (A) has no parent node but has child nodes representing downstream consequential gene modules connected to it (in blue with red edges). In turn, the sink node (B) has no child nodes but has parent nodes representing upstream causal gene modules connected to it (in blue with red edges). Note that the M04 and M07 nodes are associated with both source and sink nodes. The thickness of the edges corresponds to the number of times (in percent) the edges were detected in the 1000 permutations used to generate the Bayesian network. (C) Characterization of source- and sink-associated nodes using gene ontology (GO) and cell type (from Figure 2). Enriched GO terms are shown on the left, with significance shown as a bar graph on the right. Enrichment of cell-type clusters (top labels, same as Figure 2C) in genes of up- and downstream nodes. The lighter-to-darker shades of pink indicate increasing significance of cell-type enrichment.

Fifty consensus modules were identified that contained 21–1,575 genes. Of these, 33 were significantly enriched for at least one GO functional category (GOBP, GOMF, or GOCC) at FDR < 0.05 (Table S6) and were used to create the source and sink DAGs (Figure 4A, B). The source node (Figure 4A) served as the root of the source DAG and, in the order of the association probability (lowest to highest), were parent to the following nodes: M05, enriched (Figure 4C, bar graphs) with *membrane-bound organelle*; M10, enriched with *extracellular matrix*; M12, enriched with *cellular response to unfolded protein*; M47, enriched with *transcription regulator activity*; M04, enriched with *regulation of developmental process*; and M07, enriched with *GABAergic neuron differentiation*. The sink node (Figure 4B) was connected to two parent nodes (M04 and M07), which also served as child nodes in the source DAG (Figure 4A). In both DAGs, the M04 and M07 nodes had a parent and child relationship, respectively.

We then looked for the enrichment of the discriminant markers of 21 (Figure S2, Table S4) cell- type clusters in the up- and downstream nodes (Figure 4C, boxes). The M04 and M07 nodes were found in both the source and sink DAGs; M04 was enriched in neurogenesis and endothelial cells and polydendrocytes, while M07 was enriched in all interneurons and neurogenesis cells. Among the nodes that were associated exclusively with the source DAG, M10 was enriched in mural and endothelial cells, M01 was enriched in microglia and endothelial cells, M47 was enriched in CA3 pyramidal neurons and endothelial cells, and M05 was enriched in CA1|CA2 pyramidal neurons. Node M12 was not enriched in any of the cell types. To determine if the source- and sink-associated nodes are predictive of the epileptic phenotype, we fitted a decision tree classifier (Agrahari et al., 2018) to the eigengene representing these nodes (Table S6). Node M07 predicted the withheld control and epilepsy samples with 98% accuracy. Furthermore, the genes of node M07 were significantly overlapped with predictive genes (P + 5.46 × 10^−3^) and gene sets involving *epileptic seizures* (P + 7.89 × 10^−4^), *clonic seizures* (P + 2.84 × 10^−3^), and *myokymia with neonatal epilepsy* (*q* = 7.91 × 10^−3^).

Taken together, these results showed that the Bayesian network organized the biological events observed across control and epileptic samples into a coherent and directional graph. While these events occurred in the context of TLE, the direct association of the source and sink nodes with seven DAG nodes suggests that cell communication, GABA neuron differentiation, *extracellular matrix*, and transcription regulation are more proximal and potentially causal or consequential of TLE. Moreover, the cell-enrichment analyses of nodes M04 and M07, which were found both in the source and sink DAGs, implicate that the interaction between non-neuronal cells and GABAergic interneurons are potential mediators of the identified causal and consequential events in TLE.

### Known epilepsy drugs and related targets are associated with the source- and sink-associated nodes

We hypothesized that, if the up- and downstream nodes are directly associated with the epileptic state, drugs that mimic (Figure 5, bottom panel) or antagonize (Figure 5, top panel) their gene expression profiles should have epilepsy-inducing or antiepileptic effects, respectively. Therefore, we probed the gene sets associated with the up- and downstream nodes (Table S7) against the connectivity map, a database cataloging signatures of known drugs in diverse cell lines (Lamb, 2007).

**Figure 5:**
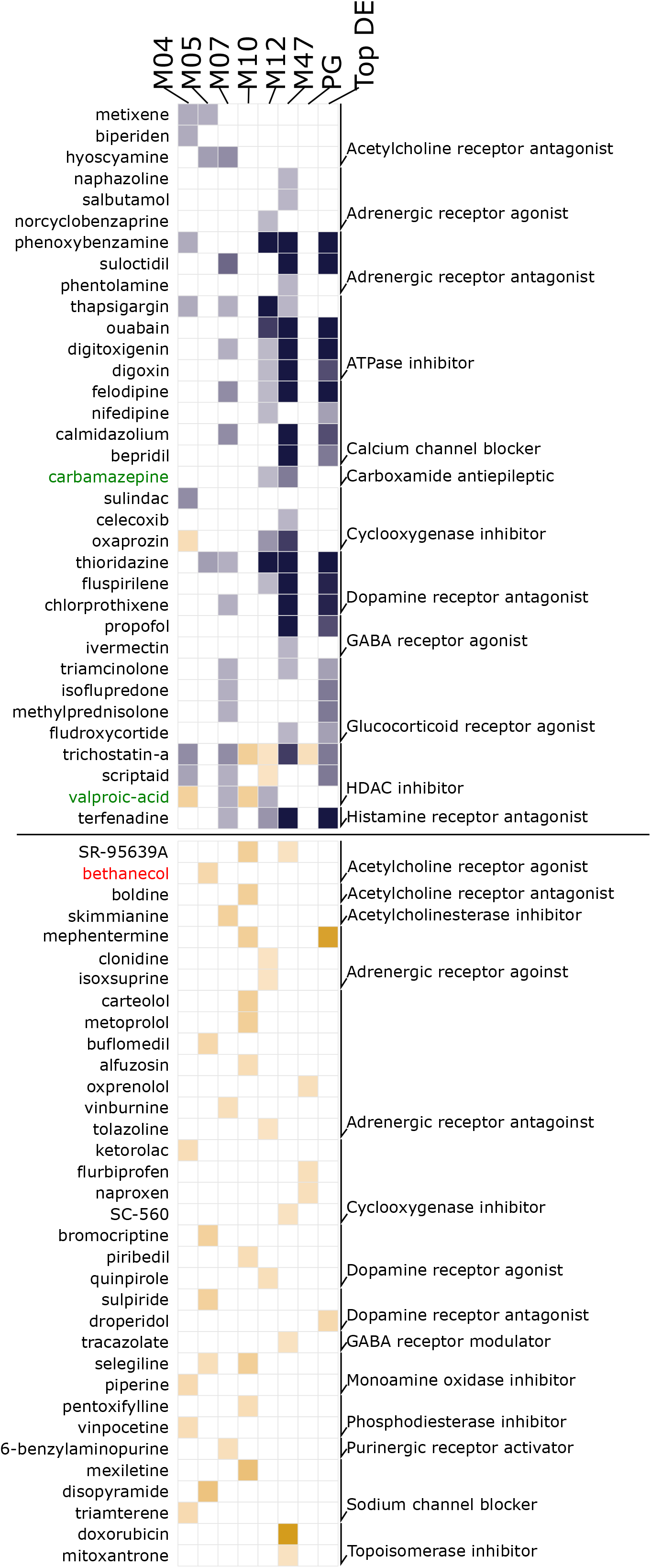
Drugs antagonizing or mimicking the upstream and downstream nodes associated with gene expression profiles. Gene sets of nodes (WGCNA modules) directly connected to source and sink nodes (M04, M05, M07, M10, M12, and M47), predictive gene lists, and top differentially expressed genes (Top DEGs) were used to probe drug-associated signatures from the connectivity map database. Note that known antiepileptic (*Carbamazepine* and *Valproic acid*, in green) and epilepsy-inducing (*bethanecol*, in red) drugs are enriched in nodes directly connected to source and sink nodes, but not in the predictive or top differentially expressed genes. Blue: therapeutic drugs; orange: pro-disease drugs. The -log_10_ of the enrichment q-value is plotted. The lighter-to-darker shades of blue and orange represent a lower-to-higher enrichment, respectively. See details in the Methods and Tables S7 and S8.

Across all seven source- and sink-associated nodes (Figure 4A, B), 59 mimicking and 212 antagonizing drugs with various modes of action were identified (*q* < 0.05, Figure 5, Table S4). Consistent with the known effects of *pilocarpine* (an acetylcholine receptor agonist) for inducing TLE (Curia et al., 2008), *SR-95639A* and *bethanechol* (Turski et al., 1983), which are drugs with a similar mode of action to that of pilocarpine, were observed as epilepsy-inducing drugs. Similarly, the antiepileptic drugs *carbamazepine* (Cereghino et al., 1974) and *valproic acid* (Romoli et al., 2018) were observed as epilepsy-antagonizing drugs. *Carbamazepine* was enriched in the M47 node, associated with pyramidal neurons (Figure 4B); *valproic acid* was enriched in the M07 node, associated with interneurons; and *carbamazepine* and *valproic acid* were enriched in module M12, which was not associated with any cell type (Figure 4B). Interestingly, valproic acid had an epilepsy-inducing effect in node M04, associated with oligodendrocytes, and node M10, associated with polydendrocytes, suggesting a diverse mechanism of action across various cell types. Notably, these drugs were not enriched in gene sets associated with predictive genes (Figure 5, column 7) and DEGs (Figure 5, column 8), which is commonly used in drug repurposing approaches (Lamb et al., 2006), indicating that the up- and downstream nodes are linked to the epileptic state.

Regarding drug targets (Table S8), those involved in transcription (*topoisomerase*), neurotransmission (*GABA receptor, norepinephrine reuptake, acetylcholine receptor, dopamine receptor*, and *monoamine oxidase*), ion balance (*L-type calcium channel and sodium channel*), hormonal balance (*estrogen receptor*), and signal transduction (*mTOR, TP53, and NFkB pathway*) were common to epilepsy-inducing and antiepileptic drugs, suggesting that these targets serve as major nodes regulating TLE. The mode of action (and target) unique to epilepsy-inducing effects involved nuclear receptor agonists (*retinoid receptor* and *constitutive androstane receptor*), metabolic control inhibitors (*thyroid peroxidase, antifolate, dehydrogenase, and carbonic anhydrase*), processes involving inhibition of immunomodulation (*atherogenesis and cytokine production*), and inhibition of signal transduction (*PKC* and *PKA*). Similarly, unique targets to antagonize epilepsy involved homeostasis through hormonal balance (*estrogen, steroid, corticosteroid, prolactin, insulin, progestogen, and thyroid*), inhibition of cellular processes (*protein synthesis, TNF production, DNA replication, DNA synthesis, RNA synthesis, dopamine uptake*, and *cell proliferation*), and inhibition of enzymes with proliferative and detoxification activity. Some known medications (psychoactive, laxative, local anesthetic, antiarrhythmic, SSRI, anti-inflammatory, tricyclic antidepressant, and thiazide diuretic drugs) also exhibited a potential epilepsy-antagonizing effect.

Together, the drug-based perturbation studies validate our Bayesian inference, and the associated mode of action suggests that the induction of epilepsy involves the facilitation of gene expression (via nuclear receptors), the inhibition of immune modulation, and metabolic control, whereas the therapeutic effects mostly involve suppression of these processes.

## DISCUSSION

Unraveling the complex biological disturbances in epilepsy is a major challenge. By integrating the transcriptomics datasets available for pilocarpine-induced TLE and single-cell RNAseq data from the mouse hippocampus, we examined the complex phenotype of TLE at multiple levels. First, we showed that there were obvious differences between the functionalities associated with down- and upregulated genes (Figure 1C). The downregulated genes were mostly associated with altered cell–cell *signaling*, whereas the upregulated genes were associated with increased homeostasis, processes involving the *extracellular matrix*, and immune and defense responses. We also showed that the subtle-to-moderate changes in gene expression associated with astrocytes and pyramidal cells of the dentate gyrus and hippocampal subfields can predict most forms of epilepsy. Second, using deconvolution analyses, we showed that different cell types exhibited disparate changes in cell proportion (Figure 2C) and that the overall functional alteration had different cellular effects (Figure 3B). Third, using a Bayesian network of the modules characterized for pathways and cell types (Figure 4), we identified several potential upstream (causal) and downstream (consequential) mechanisms associated with TLE. Furthermore, as suggested by the enrichment of markers associated with seizure and clonic seizures, these nodes, unlike the predictive gene sets (representing familial forms of epilepsy), represented status epilepticus. Finally, in a separate *in silico* analysis using gene expression profiles, we showed that drugs with both epilepsy-inducing and -antagonizing effects were associated with causal and consequential nodes (Figure 5). Collectively, our integrative transcriptomics approach provided novel insights into the cellular pathologies associated with TLE and streamlined a drug discovery/repositioning approach targeting the probable TLE causing nodes.

We observed TLE-specific changes at two levels. At the macro scale, the sparse-classifier-based predictive gene set revealed subtle changes enriched in the astrocyte, dentate gyrus, and hippocampal subfields. Notably, these changes were enriched in various forms of epilepsy (but not seizures) and suggest the involvement of a canonical structural substrate in TLE. Although the involvement of these hippocampal structures has been confirmed (Blümcke et al., 2013; Kim et al., 2015), further support for this finding stemmed from the deconvolution analysis, i.e., highly variable cell proportions associated with CA subfield neurons. This is indicative of variable cell states (De Jong et al., 2019), possibly through the transcriptional regulation (Rinotta et al., 2011) observed in the highest log-fold change bin (Figure 1C). Notably, although the deconvolution analysis suggests variable proportions of cell types, an increased proportion of neurons, which are postmitotic cells that seldom divide, is questionable. However, the increased cell proportion may be associated with other cell-level covariates that regulate gene expression, including cell size and activity (Zaitsev et al., 2019), which were not considered in our analysis. Furthermore, deconvolution analysis, consistent with the known loss of interneurons in TLE (Dudek and Shao, 2003), revealed a reduced proportion of *Sst-* and *Pvalb*-positive interneurons. However, *Vip*-positive interneurons were increased (Figure 2C), suggesting a role for these interneurons in increasing the epileptic excitotoxicity via its disinhibitory function (Pi et al., 2013). *Vip*-positive interneurons selectively receive acetylcholine and serotonergic afferents (Chédotal et al., 1994; Eckenstein and Baughman, 1984) and play a role in the long-term dilation of blood micro vessels (Cauli et al., 2004). Thus, increased activity of these neurons is suggestive of a homeostatic vascular dilation triggered by distant input events.

At the neural circuit level, the Bayesian network, which indicated the triggering of seizures via diverse mechanisms, enumerated many potential upstream causal nodes. However, node M04, which was enriched in components of the blood–brain barrier (endothelial cells, oligodendrocytes, and neurogenesis cells) (Hamanaka et al., 2018; Maki, 2017) and node M07, which was enriched in interneurons, were identified as both causal and consequential nodes, possibly suggesting an activity (seizure)-dependent oscillation between the node elements, leading to seizure amplification. Furthermore, a perhaps stronger support for causal associates of nodes M04 and M07 stemmed from the significant enrichment of valproic acid, a potent AED (Romoli et al., 2018), in the genes associated with these nodes (Figure 5). While more comprehensive and experimental studies are required to understand the cross talk between interneurons (M07) and non-neuronal cells (M04) in the seizure phenotype, the interneuron-specific preference of oligodendrocytes (Zonouzi et al., 2019), activity-dependent oscillatory behavior of oligodendrocytes (Pajevic et al., 2014), seizure-like activity in impaired endothelial cells (Li et al., 2018), antiepileptic role of histone deacetylase inhibitors targeting oligodendrogenesis (Gibson et al., 2014), and accelerated myelination (function of oligodendrocytes) in the promotion of seizures (Goldsberry et al., 2011) are known. Accordingly, our analysis revealed two crucial elements in TLE: a canonical structure involved in TLE possibly representing the epileptic focus; and a microcircuitry node involving elements of the BBB and interneurons in the potential amplification of seizures.

Causal associations are usually validated using perturbation studies (Meinshausen et al., 2016). As these critical data were lacking, a final set of *in silico* analyses were conducted to utilize the gene expression signatures from drug perturbation databases. While validating our Bayesian inference, we observed that known drugs with epilepsy-inducing or epilepsy-antagonizing effects were associated with causal and consequential nodes. Furthermore, on examining the mode of action associated with each drug, we differentiated the key nodes defined by their potential to trigger an epilepsy-inducing or epilepsy-reversing effect. Interestingly, together with the aforementioned biological processes (transcription regulation (Morgan and Curran, 1991) and distant serotonergic and cholinergic inputs (Zhan et al., 2016)), we observed other processes involving hormonal balance (Velíšková and DeSantis, 2013) and signal transduction through mTOR (Crino, 2015) and L-type sodium channels (Stafstrom, 2007) reported in other forms of epilepsy. Finally, the modes of action associated with therapeutic drugs were mostly associated with the inhibition of homeostatic, proliferative, and detoxification processes, suggesting that TLE largely comprises maladaptive processes. Therefore, the aforementioned increased activity of *Vip*-positive interneurons suggests a maladaptive dilation of the vascular system, which could initially be triggered to maintain blood flow to meet the TLE-associated increased metabolic demands.

This study was performed using well-powered, publicly available datasets to study the widest possible biological events associated with TLE rather than the specificities associated with sex and age. The results suggesting cellular correlates and causal and consequential links were validated using *in silico* approaches; however, they should be interpreted as hypothesis-generating findings for subsequent experimental confirmation.

## MATERIALS AND METHODS

### Data

This study used data previously collected by (Srivastava et al., 2018), who performed high-throughput RNA sequencing on whole hippocampus samples from pilocarpine-treated (n + 108) and littermate control (n + 103) adult (>12 weeks) male Crl:NMRI(Han)-FR mice. Pilocarpine-treated mice developed convulsive seizures and effectively modeled status epilepticus (Srivastava et al., 2018). Fastq.gz files for all samples were downloaded from the European Nucleotide Archive (accession number: PRJEB18790) and were aligned to the mouse reference genome GRCm38 (provided by Ensembl) using HISAT2 aligner. Count data were generated for the reads aligned to exons and transcripts using GenomicFeature and GenomicAlignments (Lawrence et al., 2013) in R and the mouse gene model (GTF file) provided by Ensembl.

### Differential expression analysis

After filtering out genes present in a minimum of 100 samples, 15,300 genes (Table S1) were examined for differential expression between the control and epileptic groups using Deseq2 in R. To segregate the effects of subtle and obvious changes, all differentially expressed genes (DEGs) (false discovery rate [FDR]-corrected *P*-value < 0.05) were binned based on their log-fold change (Figure 1B). To filter out the minimal set of genes that could discriminate the epileptic phenotype from the control phenotype, we used a sparse classification strategy employing nearest shrunken centroids (Tibshirani et al., 2002) and the diagonal discriminant classifiers (Dudoit et al., 2002) implemented by MLseq in R (Zararsiz et al., 2017). The classification model identified 119 genes (termed predictive genes, Table S3) that were segregated as being up- or downregulated based on their initial Deseq2-based log-fold change (Table S1). To characterize the predictive genes, we searched for their enrichment in gene ontology (GO) terms, different log-fold change bins of DEGs, and cell type-specific discriminant markers. In all enrichment analyses, we used the hypergeometric overlap analysis implemented by GeneOverlap in R. A background of 21196 genes and a significant cut-off q-value < 0.05 were used.

### Pathway enrichment analysis

To identify the biological pathways affected at the different log-fold change bins of DEGs, we searched for the enrichment of different GO terms in the Biological Pathway (GOBP), Molecular Function (GOMF), and Cellular Component (GOCC) categories using the GO database. To reduce and catalog our long list of GO terms into an interpretable format, we clustered the pathways based on biological themes (Shukla et al., 2020b, 2018). Briefly, the theme for a given pathway was selected based on either a text search, in which the name of the theme was used as a keyword for text query, or on the parent–child association between the GO terms in our list of significant pathways (child pathways) and the handpicked pathways in the GO database (parent pathway) representing the theme using GOdb in R.

### Cellular deconvolution analysis

To estimate the TLE-associated changes in cell type proportion and the genes and pathways affected by an altered cell type proportion, we employed the previously described deconvolution analysis (Baron et al., 2016; Aaron M. Newman et al., 2015; Shen-Orr et al., 2010; Shukla et al., 2020b) in the bscqsc package in R. The analysis involved four steps. A) Identifying cell type-specific marker genes: using the single-cell RNAseq dataset for mouse hippocampus available from the Gene Expression Omnibus (accession: GSE116470) (Saunders et al., 2018), we identisfied clusters of cell types (Figure 2A) and markers that were specific to each cluster (Figure S2, Table S4) using SEURAT in R for single-cell analysis (Stuart et al., 2019). To segregate cell clusters based on subtle differences in expression, the resolution parameter was set to 1.2. B) Building the reference signature matrix of marker genes (Figure S2): this matrix contained the highly discriminatory marker gene expression of each cell-type cluster averaged across all cells of a given cluster. A gene was considered cluster-specific only when it showed a ³3-fold difference in expression compared with that in other cell-type clusters. C) Estimating proportions: the resulting reference signature matrix was used to estimate the cluster-specific cell type proportion in control and epileptic samples using Support Vector Regression implemented by CIBERSORT in R (Aaron M Newman et al., 2015). The Wilcoxon test was used to estimate the significance of altered cell-type proportions. D) Adjusting the bulk tissue gene expression for differences in cell-type proportion: to assess the effect of altered proportions of different cell types, we statistically regressed out the effect of cell-type clusters representing pyramidal neurons, interneurons, and non-neuronal cells that showed significant changes in proportion in TLE vs. control samples. This was implemented by expanding the design model used for examining the differential expression between TLE and control subjects by incorporating the estimated cell-type proportion (from step C) in the design matrix. Genes with further reduced *P*-values (Table S5) were considered as being affected by the altered cell-type proportion and were applied to a pathway analysis (Figure 3B) using the GO database.

### Bayesian network analysis

To identify the probabilistic upstream causal and downstream consequential associations between TLE and gene coexpression modules representing different biological themes, we used a weighted gene coexpression network analysis (WGCNA) (Zhao et al., 2010) and Pigengene (Foroushani et al., 2017) in R. For each cohort, count data that were normalized based on the size factor (using DESeq2 in R) were used to construct a matrix of pairwise Pearson’s correlations between genes, which was then transformed to a signed adjacency matrix using a power of β + 12. Adjacency matrices provide the connectedness between genes. To calculate the interconnectedness between genes (which defines “modules”), we derived the topological overlap, a biologically meaningful measure of the similarity between two genes based on their coexpression relationship with all other genes (Yip and Horvath, 2006). Consensus modules between the control and epilepsy cohorts were then identified using the blockwiseConsensusModules function. Fifty consensus modules numerically labeled 1–50, were functionally characterized using the GO database for the GOBP, GOMF, and GOCC GO categories; the top representative pathways with a Bonferroni-corrected *P*-value of <0.05 (Table S6) were used to label the modules. Modules that were not characterized and module 0, which comprised genes not assigned to any module (Langfelder and Horvath, 2008), were not analyzed further.

The first principal component (the eigengene) summarized a given module by accounting for the maximum variability of all its constituent genes (Langfelder and Horvath, 2007) and was used as a biological signature (Agrahari et al., 2018) of the module and to identify the mechanisms associated with TLE. We used Pigengene in R, which works on this principle using the module eigengenes as random variables to train a Bayesian network by modeling the probabilistic dependencies between all modules as directed acyclic graphs (DAGs, Figure 4A, B; modules in the DAG network are termed “nodes”). In addition, the network had a categorical variable that modeled the disease state. Its value corresponded to the labels associated with the samples, i.e., control and epilepsy. To simplify the inference, the disease node served either as a source (with no parent terms above it) or as a sink (with no child terms below it) (Foroushani et al., 2017; Shukla et al., 2020b). This was implemented by blacklisting all incoming and outgoing edges to the source and node, respectively. To search for the optimal Bayesian network fitting our data, we performed 1000 permutations using the default parameters in Pigengen.

### Drug target enrichment analysis

To identify causal/consequential nodes associated with epilepsy-inducing and therapeutic drugs, we first segregated the up- and downregulated genes (Table S7) associated with the nodes (WGCNA modules) that were directly connected to source and sink nodes (Figure 4 A and B, nodes in blue) based on their log-fold change estimates derived from the differential expression analysis (Table S1). Next, drug-specific gene markers (associated with the connectivity map database (Lamb, 2007)) that were attributed to up- and downregulated effects were downloaded from the Enricher library of gene sets (Kuleshov et al., 2016), and significant enrichment of drug-induced molecular signatures in the gene sets associated with causal and consequential nodes (Table S7) were calculated using hypergeometric overlap analysis. Based on the signature-reversing principle (Shukla et al., 2020a), an enrichment in the inverse direction (i.e., up- or downregulated drug signature enriched in down- or upregulated WGCNA module genes, respectively) was considered therapeutic, whereas an enrichment in the similar direction (i.e., up- or downregulated drug signature enriched in up- or downregulated WGCNA module genes, respectively) was considered disease inducing. Gene markers of drugs with known modes of action and targets were used to understand druggable mechanisms (Table S8) and targets.

## DATA AVAILABILITY

The datasets used in this study are available in the following databases:

- RNA-Seq data: European Nucleotide Archive (ENA; http://www.ebi.ac.uk/ena) under accession number PRJEB18790.
- Single-cell RNA-seq data: Gene Expression Omnibus https://www.ncbi.nlm.nih.gov/geo/query/acc.cgi?acc+GSE116470

## AUTHOR CONTRIBUTIONS

R.S. and N.D.H. conceptualized the study, performed the analysis and together wrote the manuscript. N.D.H., M.A.S., X.W., and H.M.E. participated in gene-ontology analysis. H.A.E. and N.F. participated in deconvolution and Bayesian-network analysis. P.R.C. and R.E.M. participated in validating the causal inference using cmap database.

---

**Figure S1a:**
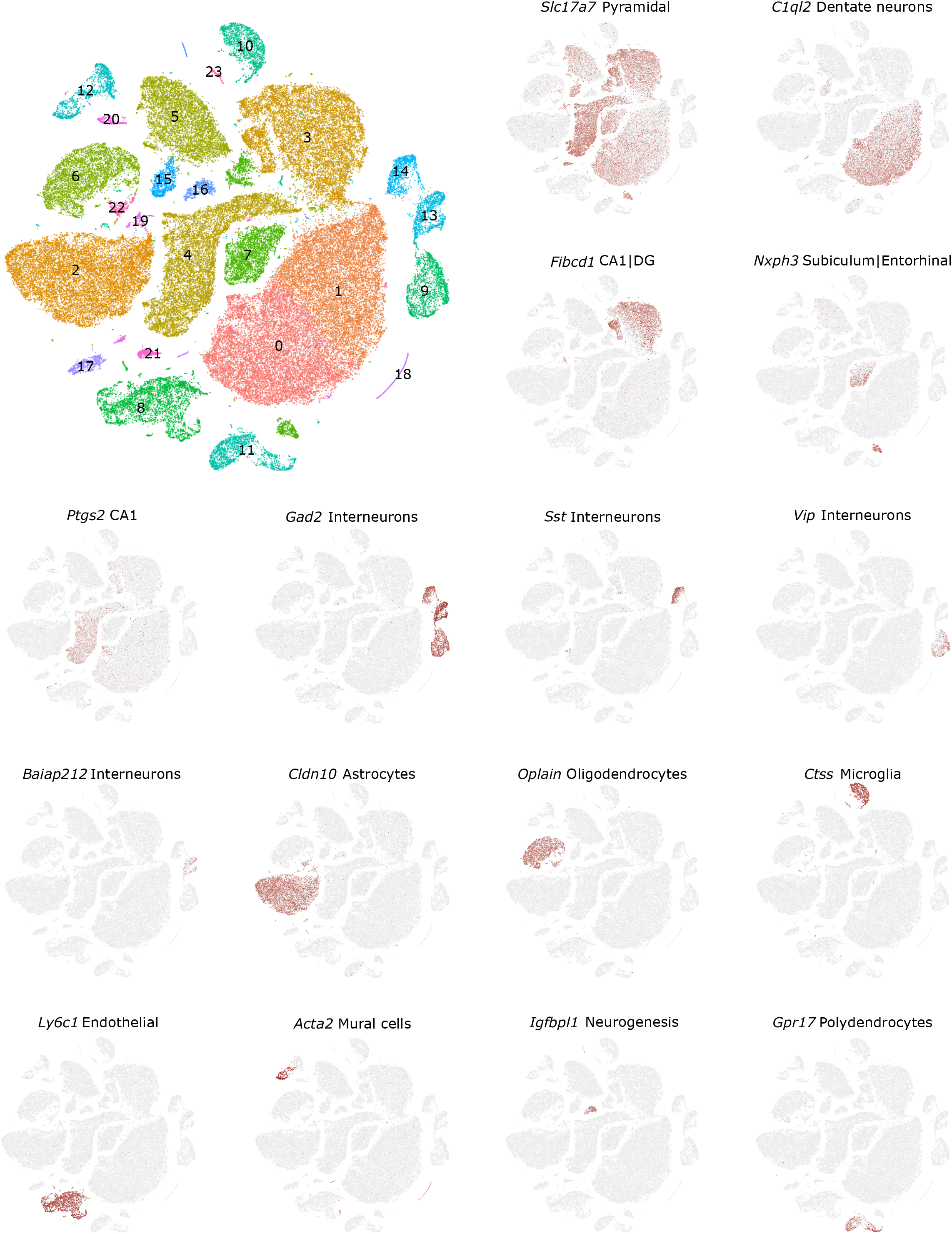

**Figure S1b:**
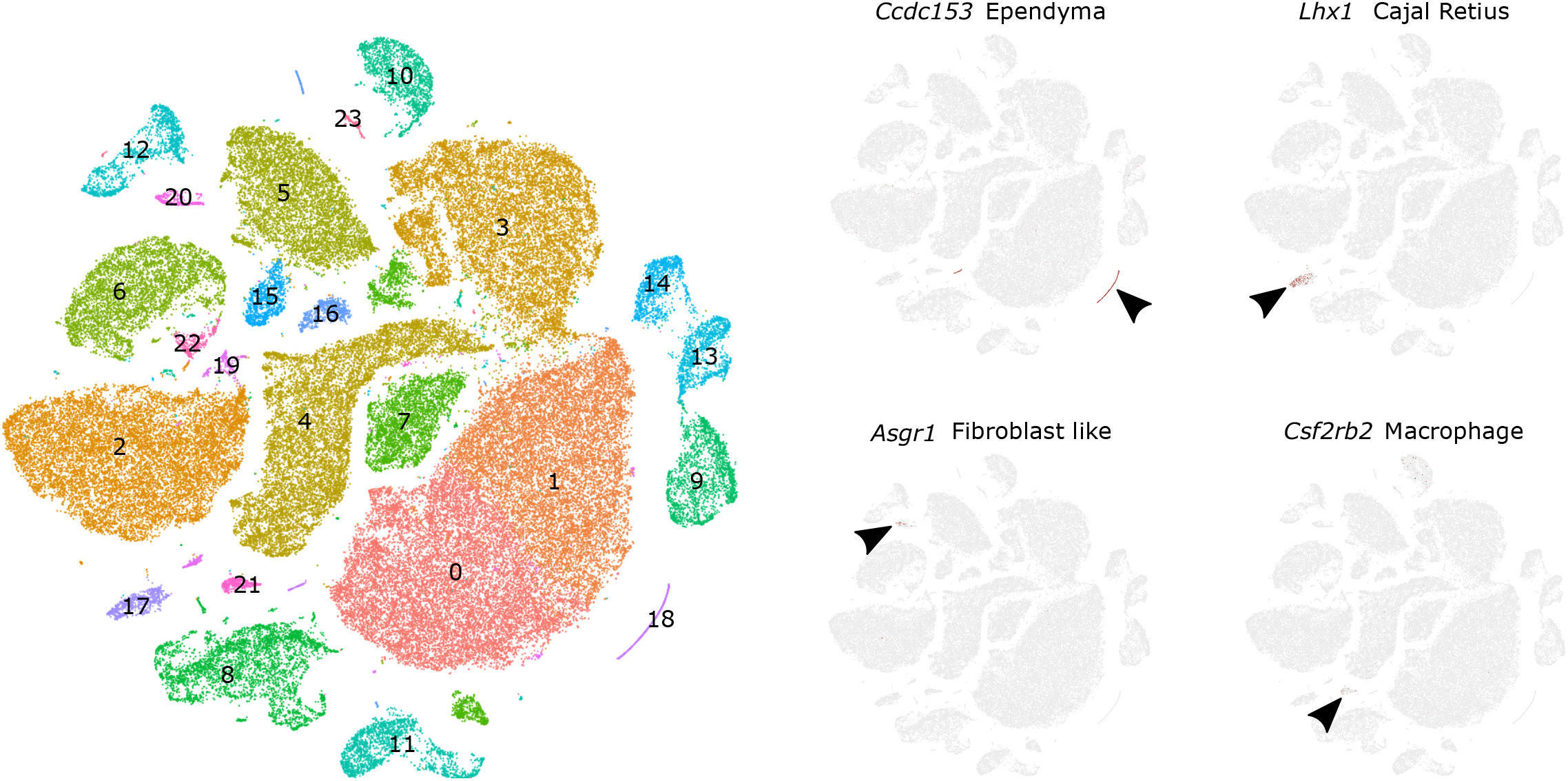

**Figure S2:**
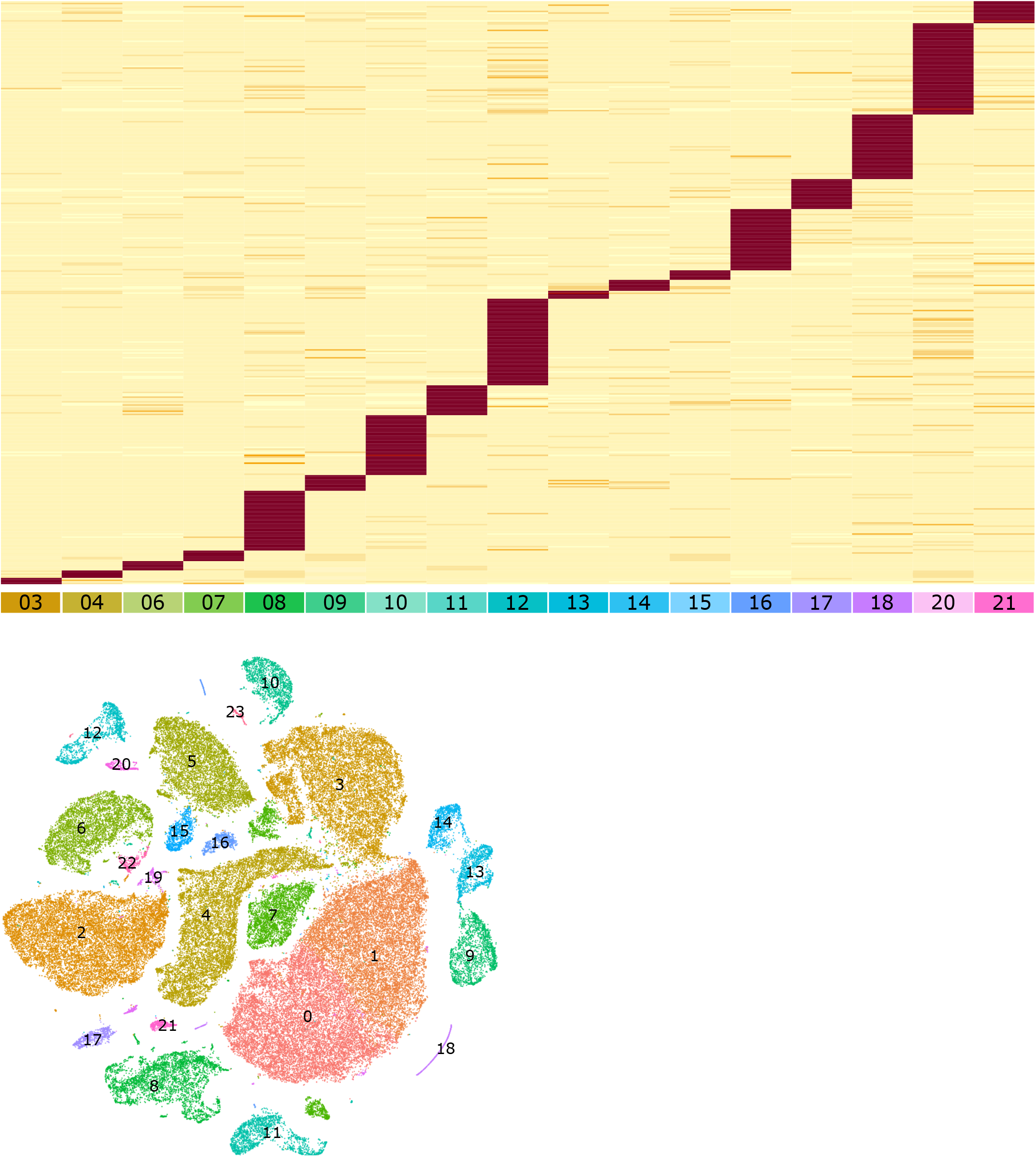

## REFERENCES

Agrahari R, Foroushani A, Docking TR, Chang L, Duns G, Hudoba M, Karsan A, Zare H. 2018. Applications of Bayesian network models in predicting types of hematological malignancies. Sci Rep. doi:10.1038/s41598-018-24758-5

Baron M, Veres A, Wolock SL, Faust AL, Gaujoux R, Vetere A, Ryu JH, Wagner BK, Shen-Orr SS, Klein AM, Melton DA, Yanai I. 2016. A Single-Cell Transcriptomic Map of the Human and Mouse Pancreas Reveals Inter- and Intra-cell Population Structure. Cell Syst. doi:10.1016/j.cels.2016.08.011

Berkovic SF, Mulley JC, Scheffer IE, Petrou S. 2006. Human epilepsies: interaction of genetic and acquired factors. Trends Neurosci. doi:10.1016/j.tins.2006.05.009

Blair RDG. 2012. Temporal Lobe Epilepsy Semiology. Epilepsy Res Treat. doi:10.1155/2012/751510

Blümcke I, Thom M, Aronica E, Armstrong DD, Bartolomei F, Bernasconi A, Bernasconi N, Bien CG, Cendes F, Coras R, Cross JH, Jacques TS, Kahane P, Mathern GW, Miyata H, Moshé SL, Oz B, Özkara Ç, Perucca E, Sisodiya S, Wiebe S, Spreafico R. 2013. International consensus classification of hippocampal sclerosis in temporal lobe epilepsy: A Task Force report from the ILAE Commission on Diagnostic Methods. Epilepsia. doi:10.1111/epi.12220

Cauli B, Tong XK, Rancillac A, Serluca N, Lambolez B, Rossier J, Hamel E. 2004. Cortical GABA interneurons in neurovascular coupling: Relays for subcortical vasoactive pathways. J Neurosci. doi:10.1523/JNEUROSCI.3065-04.2004

Cereghino JJ, Brock JT, van Meter JC, Klffin Penry J, Smith LD, White BG. 1974. Carbamazepine for epilepsy: A controlled prospective evaluation. Neurology. doi:10.1212/wnl.24.5.401

Chédotal A, Cozzani C, Faure MP, Hartman BK, Hamel E. 1994. Distinct choline acetyltransferase (ChAT) and vasoactive intestinal polypeptide (VIP) bipolar neurons project to local blood vessels in the rat cerebral cortex. Brain Res. doi:10.1016/0006-8993(94)90076-0

Christensen J, Pedersen MG, Pedersen CB, Sidenius P, Olsen J, Vestergaard M. 2009. Long-term risk of epilepsy after traumatic brain injury in children and young adults: a population-based cohort study. Lancet. doi:10.1016/S0140-6736(09)60214-2

Crino PB. 2015. mTOR *signaling* in epilepsy: Insights from malformations of cortical development. Cold Spring Harb Perspect Med. doi:10.1101/cshperspect.a022442

Curia G, Longo D, Biagini G, Jones RSG, Avoli M. 2008. The pilocarpine model of temporal lobe epilepsy. J Neurosci Methods. doi:10.1016/j.jneumeth.2008.04.019

De Jong T V., Moshkin YM, Guryev V. 2019. Gene expression variability: The other dimension in transcriptome analysis. Physiol Genomics. doi:10.1152/physiolgenomics.00128.2018

DeFelipe J. 1999. Chandelier cells and epilepsy. Brain. doi:10.1093/brain/122.10.1807

Dixit AB, Banerjee J, Srivastava A, Tripathi M, Sarkar C, Kakkar A, Chandra PS. 2016. RNA-Seq analysis of hippocampal tissues reveals novel candidate genes for drug refractory epilepsy in patients with MTLE-HS. Int J Epilepsy. doi:10.1016/j.ijep.2015.12.006

Dudek FE, Shao L-R. 2003. Loss of GABAergic Interneurons in Seizure-Induced Epileptogenesis. Epilepsy Curr. doi:10.1046/j.1535-7597.2003.03503.x

Dudoit S, Fridlyand J, Speed TP. 2002. Comparison of discrimination methods for the classification of tumors using gene expression data. J Am Stat Assoc. doi:10.1198/016214502753479248

Eckenstein F, Baughman RW. 1984. Two types of cholinergic innervation in cortex, one co-localized with vasoactive intestinal polypeptide. Nature. doi:10.1038/309153a0

Foroushani A, Agrahari R, Docking R, Chang L, Duns G, Hudoba M, Karsan A, Zare H. 2017. Large-scale gene network analysis reveals the significance of *extracellular matrix* pathway and homeobox genes in acute myeloid leukemia: An introduction to the Pigengene package and its applications. BMC Med Genomics. doi:10.1186/s12920-017-0253-6

Frishberg A, Peshes-Yaloz N, Cohn O, Rosentul D, Steuerman Y, Valadarsky L, Yankovitz G, Mandelboim M, Iraqi FA, Amit I, Mayo L, Bacharach E, Gat-Viks I. 2019. Cell composition analysis of bulk genomics using single-cell data. Nat Methods. doi:10.1038/s41592-019-0355-5

Gibson EM, Purger D, Mount CW, Goldstein AK, Lin GL, Wood LS, Inema I, Miller SE, Bieri G, Zuchero JB, Barres BA, Woo PJ, Vogel H, Monje M. 2014. Neuronal activity promotes oligodendrogenesis and adaptive myelination in the mammalian brain. Science (80-). doi:10.1126/science.1252304

Goldsberry G, Mitra D, MacDonald D, Patay Z. 2011. Accelerated myelination with motor system involvement in a neonate with immediate postnatal onset of seizures and hemimegalencephaly. Epilepsy Behav. doi:10.1016/j.yebeh.2011.06.025

Guerrini R, Casari G, Marini C. 2003. The genetic and molecular basis of epilepsy. Trends Mol Med. doi:10.1016/S1471-4914(03)00116-3

Hamanaka G, Ohtomo R, Takase H, Lok J, Arai K. 2018. White-matter repair: Interaction between oligodendrocytes and the neurovascular unit. Brain Circ. doi:10.4103/bc.bc_15_18

Jiruska P, Csicsvari J, Powell AD, Fox JE, Chang WC, Vreugdenhil M, Li X, Palus M, Bujan AF, Dearden RW, Jefferys JGR. 2010. High-frequency network activity, global increase in neuronal activity, and synchrony expansion precede *epileptic seizures* in vitro. J Neurosci. doi:10.1523/JNEUROSCI.0535-10.2010

Kim J Bin, Suh S il, Kim JH. 2015. Volumetric and shape analysis of hippocampal subfields in unilateral mesial temporal lobe epilepsy with hippocampal atrophy. Epilepsy Res. doi:10.1016/j.eplepsyres.2015.09.004

Kuleshov M V., Jones MR, Rouillard AD, Fernandez NF, Duan Q, Wang Z, Koplev S, Jenkins SL, Jagodnik KM, Lachmann A, McDermott MG, Monteiro CD, Gundersen GW, Ma’ayan A. 2016. Enrichr: a comprehensive gene set enrichment analysis web server 2016 update. Nucleic Acids Res. doi:10.1093/nar/gkw377

Lamb J. 2007. The Connectivity Map: A new tool for biomedical research. Nat Rev Cancer. doi:10.1038/nrc2044

Lamb J, Crawford ED, Peck D, Modell JW, Blat IC, Wrobel MJ, Lerner J, Brunet J-P, Subramanian A, Ross KN, Reich M, Hieronymus H, Wei G, Armstrong SA, Haggarty SJ, Clemons PA, Wei R, Carr SA, Lander ES, Golub TR. 2006. The Connectivity Map: using gene-expression signatures to connect small molecules, genes, and disease. Science 313:1929–35. doi:10.1126/science.1132939

Langfelder P, Horvath S. 2008. WGCNA: an R package for weighted correlation network analysis. BMC Bioinformatics 9:559. doi:10.1186/1471-2105-9-559

Langfelder P, Horvath S. 2007. Eigengene networks for studying the relationships between co-expression modules. BMC Syst Biol 1:54. doi:10.1186/1752-0509-1-54

Lawrence M, Huber W, Pagès H, Aboyoun P, Carlson M, Gentleman R, Morgan MT, Carey VJ. 2013. Software for Computing and Annotating Genomic Ranges. PLoS Comput Biol. doi:10.1371/journal.pcbi.1003118

Lévesque M, Avoli M, Bernard C. 2016. Animal models of temporal lobe epilepsy following systemic chemoconvulsant administration. J Neurosci Methods. doi:10.1016/j.jneumeth.2015.03.009

Li S, Kumar P, Joshee S, Kirschstein T, Subburaju S, Khalili JS, Kloepper J, Du C, Elkhal A, Szabó G, Jain RK, Köhling R, Vasudevan A. 2018. Endothelial cell-derived GABA *signaling* modulates neuronal migration and postnatal behavior. Cell Res. doi:10.1038/cr.2017.135

Löscher W. 2011. Critical review of current animal models of seizures and epilepsy used in the discovery and development of new antiepileptic drugs. Seizure. doi:10.1016/j.seizure.2011.01.003

Maki T. 2017. Novel roles of oligodendrocyte precursor cells in the developing and damaged brain. Clin Exp Neuroimmunol. doi:10.1111/cen3.12358

Mancarci BO, Toker L, Tripathy SJ, Li B, Rocco B, Sibille E, Pavlidis P. 2017. Cross-laboratory analysis of brain cell type transcriptomes with applications to interpretation of bulk tissue data. eNeuro. doi:10.1523/ENEURO.0212-17.2017

Meinshausen N, Hauser A, Mooij JM, Peters J, Versteeg P, Bühlmann P. 2016. Methods for causal inference from gene perturbation experiments and validation. Proc Natl Acad Sci U S A 113:7361–8. doi:10.1073/pnas.1510493113

Morgan JI, Curran T. 1991. Proto-oncogene transcription factors and epilepsy. Trends Pharmacol Sci. doi:10.1016/0165-6147(91)90594-I

Newman Aaron M., Liu CL, Green MR, Gentles AJ, Feng W, Xu Y, Hoang CD, Diehn M, Alizadeh AA. 2015. Robust enumeration of cell subsets from tissue expression profiles. Nat Methods 12:453–457. doi:10.1038/nmeth.3337

Newman Aaron M, Liu CL, Green MR, Gentles AJ, Feng W, Xu Y, Hoang CD, Diehn M, Alizadeh AA. 2015. Robust enumeration of cell subsets from tissue expression profiles. Nat Methods 12:453–457. doi:10.1038/nmeth.3337

Newman AM, Steen CB, Liu CL, Gentles AJ, Chaudhuri AA, Scherer F, Khodadoust MS, Esfahani MS, Luca BA, Steiner D, Diehn M, Alizadeh AA. 2019. Determining cell type abundance and expression from bulk tissues with digital cytometry. Nat Biotechnol. doi:10.1038/s41587-019-0114-2

Pajevic S, Basser PJ, Fields RD. 2014. Role of myelin plasticity in oscillations and synchrony of neuronal activity. Neuroscience. doi:10.1016/j.neuroscience.2013.11.007

Pfisterer U, Petukhov V, Demharter S, Meichsner J, Thompson J, Batiuk M, Asenjo Martinez A, Vasistha N, Thakur A, Mikkelsen J, Adorjan I, Pinborg L, Pers T, von Engelhardt J, Kharchenko P, Khodosevich K. 2020. Identification of epilepsy-associated neuronal subtypes and gene expression underlying epileptogenesis. Nat Commun 11:1–19. doi:10.1038/s41467-020-18752-7

Pi H-JJ, Hangya B, Kvitsiani D, Sanders JI, Huang ZJ, Kepecs A. 2013. Cortical interneurons that specialize in disinhibitory control. Nature 503:521–4. doi:10.1038/nature12676

Pitkänen A, Sutula TP. 2002. Is epilepsy a progressive disorder? Prospects for new therapeutic approaches in temporal-lobe epilepsy. Lancet Neurol. doi:10.1016/S1474-4422(02)00073-X

Rinotta R, Jaimovicha A, Friedman N. 2011. Exploring transcription regulation through cell-to-cell variability. Proc Natl Acad Sci U S A. doi:10.1073/pnas.1013148108

Romoli M, Mazzocchetti P, D’Alonzo R, Siliquini S, Rinaldi VE, Verrotti A, Calabresi P, Costa C. 2018. Valproic Acid and Epilepsy: From Molecular Mechanisms to Clinical Evidences. Curr Neuropharmacol. doi:10.2174/1570159X17666181227165722

Salzmann A, Malafosse A. 2012. Genetics of Temporal Lobe Epilepsy: A Review. Epilepsy Res Treat. doi:10.1155/2012/863702

Saunders A, Macosko EZ, Wysoker A, Goldman M, Krienen FM, de Rivera H, Bien E, Baum M, Bortolin L, Wang S, Goeva A, Nemesh J, Kamitaki N, Brumbaugh S, Kulp D, McCarroll SA. 2018. Molecular Diversity and Specializations among the Cells of the Adult Mouse Brain. Cell. doi:10.1016/j.cell.2018.07.028

Shen-Orr SS, Gaujoux R. 2013. Computational deconvolution: extracting cell type-specific information from heterogeneous samples. Curr Opin Immunol. doi:10.1016/j.coi.2013.09.015

Shen-Orr SS, Tibshirani R, Khatri P, Bodian DL, Staedtler F, Perry NM, Hastie T, Sarwal MM, Davis MM, Butte AJ. 2010. Cell type-specific gene expression differences in complex tissues. Nat Methods. doi:10.1038/nmeth.1439

Shukla R, Henkel ND, Alganem K, Hamoud A, Reigle J, Alnafisah RS, Eby HM, Imami AS, Creeden JF, Miruzzi SA, Meller J, Mccullumsmith RE. 2020a. Signature-based approaches for informed drug repurposing: targeting CNS disorders. Neuropsychopharmacology 1–15. doi:10.1038/s41386-020-0752-6

Shukla R, Prevot TD, French L, Isserlin R, Rocco BR, Banasr M, Bader GD, Sibille E. 2018. The relative contributions of cell-dependent cortical microcircuit aging to cognition and anxiety. Biol Psychiatry. doi:10.1016/J.BIOPSYCH.2018.09.019

Shukla R, Sibille E, Newton D, Sumitomo A, Zare H, Mccullumsmith R, Lewis D, Tomoda T. 2020b. Molecular Characterization of Depression Trait and State. bioRxiv 2020.04.24.058610. doi:10.1101/2020.04.24.058610

Srivastava PK, Eyll J van, Godard P, Mazzuferi M, Delahaye-Duriez A, Steenwinckel J Van, Gressens P, Danis B, Vandenplas C, Foerch P, Leclercq K, Mairet-Coello G, Cardenas A, Vanclef F, Laaniste L, Niespodziany I, Keaney J, Gasser J, Gillet G, Shkura K, Chong SA, Behmoaras J, Kadiu I, Petretto E, Kaminski RM, Johnson MR. 2018. A systems-level framework for drug discovery identifies Csf1R as an anti-epileptic drug target. Nat Commun. doi:10.1038/s41467-018-06008-4

St. Laurent G, Shtokalo D, Tackett MR, Yang Z, Vyatkin Y, Milos PM, Seilheimer B, McCaffrey TA, Kapranov P. 2013. On the importance of small changes in RNA expression. Methods. doi:10.1016/j.ymeth.2013.03.027

Stafstrom CE. 2007. Persistent Sodium Current and Its Role in Epilepsy. Epilepsy Curr. doi:10.1111/j.1535-7511.2007.00156.x

Stafstrom CE, Carmant L. 2015. Seizures and epilepsy: An overview for neuroscientists. Cold Spring Harb Perspect Biol. doi:10.1101/cshperspect.a022426

Stuart T, Butler A, Hoffman P, Hafemeister C, Papalexi E, Mauck WM, Hao Y, Stoeckius M, Smibert P, Satija R. 2019. Comprehensive Integration of Single-Cell Data. Cell. doi:10.1016/j.cell.2019.05.031

Tibshirani R, Hastie T, Narasimhan B, Chu G. 2002. Diagnosis of multiple cancer types by shrunken centroids of gene expression. Proc Natl Acad Sci U S A. doi:10.1073/pnas.082099299

Turski WA, Cavalheiro EA, Turski L, Kleinrok Z. 1983. Intrahippocampal bethanechol in rats: Behavioural, electroencephalographic and neuropathological correlates. Behav Brain Res. doi:10.1016/0166-4328(83)90026-8

Velíšková J, DeSantis KA. 2013. Sex and hormonal influences on seizures and epilepsy. Horm Behav. doi:10.1016/j.yhbeh.2012.03.018

Wang Y, Zhou D, Wang B, Li H, Chai H, Zhou Q, Zhang S, Stefan H. 2003. A kindling model of pharmacoresistant temporal lobe epilepsy in Sprague-Dawley rats induced by Coriaria lactone and its possible mechanism. Epilepsia. doi:10.1046/j.1528-1157.2003.32502.x

Yip AM, Horvath S. 2006. The Generalized Topological Overlap Matrix For Detecting Modules in Gene Networks. In BIOCOMP.

Zaitsev K, Bambouskova M, Swain A, Artyomov MN. 2019. Complete deconvolution of cellular mixtures based on linearity of transcriptional signatures. Nat Commun. doi:10.1038/s41467-019-09990-5

Zararsiz G, Goksuluk D, Klaus B, Korkmaz S, Eldem V, Karabulut E, Ozturk A. 2017. voomDDA: Discovery of diagnostic biomarkers and classification of RNA-seq data. PeerJ. doi:10.7717/peerj.3890

Zhan Q, Buchanan GF, Motelow JE, Andrews J, Vitkovskiy P, Chen WC, Serout F, Gummadavelli A, Kundishora A, Furman M, Li W, Bo X, Richerson GB, Blumenfeld H. 2016. Impaired serotonergic brainstem function during and after seizures. J Neurosci. doi:10.1523/JNEUROSCI.4331-15.2016

Zhang B, Gaiteri C, Bodea L-GG, Wang Z, McElwee J, Podtelezhnikov AAA, Zhang C, Xie T, Tran L, Dobrin R, Fluder E, Clurman B, Melquist S, Narayanan M, Suver C, Shah H, Mahajan M, Gillis T, Mysore J, MacDonald MEE, Lamb JRR, Bennett DAA, Molony C, Stone DJJ, Gudnason V, Myers AJJ, Schadt EEE, Neumann H, Zhu J, Emilsson V. 2013. Integrated systems approach identifies genetic nodes and networks in late-onset Alzheimer’s disease. Cell 153:707–720. doi:10.1016/j.cell.2013.03.030

Zhao W, Langfelder P, Fuller T, Dong J, Li A, Hovarth S. 2010. Weighted gene coexpression network analysis: State of the art. J Biopharm Stat. doi:10.1080/10543400903572753

Zonouzi M, Berger D, Jokhi V, Kedaigle A, Lichtman J, Arlotta P. 2019. Individual Oligodendrocytes Show Bias for Inhibitory Axons in the Neocortex. Cell Rep. doi:10.1016/j.celrep.2019.05.018

