## Supplementary Information for "Cellular, molecular, and therapeutic characterization of pilocarpine-induced temporal lobe epilepsy"

### **SUPPLEMENTAL FIGURES (each as pdf file)**

**Supplemental Figure S1a, S1b:** Characterizing the TSNE clusters (Figure 2A) using the cell-type specific marker genes collected from Saunders *et.al.*(Saunders et al., 2018). ***Slc17a7*:** solute carrier family 17 (sodium dependent inorganic phosphate cotransporter), member 7; **C*1ql2*:** complement component 1, q subcomponent-like 2; ***Fibcd1:*** Fibrinogen C domain containing 1; ***Nxph3:*** neurexophilin 3; ***Ptgs2*:** prostaglandin-endoperoxide synthase 2; ***Gad2*:** glutamic acid/glutamate decarboxylase 2; ***Sst*:** somatostatin; ***Vip*:** vasoactive intestinal peptide; ***Baiap*2:** brain-specific angiogenesis inhibitor 1-associated protein 2; ***Cldn10*:** claudin 10; ***Oplain*:** Transmembrane Protein 10; ***Ctss*:** cathepsin S; ***Lyc6c1:*** lymphocyte antigen 6 complex, locus C1; ***Acta2*:** actin, alpha 2, smooth muscle; ***Igflbpl1*:** insulin line growth factor protein like 1; ***Gpr1*7:** G-protein coupled receptor 17; ***Ccdc153:*** coiled-coil domain containing 153; ***Lhx1*:** LIM homeobox protein 1; ***Asgr1*:** asialoglycoprotein receptor 1; ***Csf2rb2*:** colony-stimulating factor 2, beta-2, low affinity.

**Supplemental Figure S2:** Heatmap of highly discriminative gene, i.e. markers used as signatures for estimating relative cell-type proportions in **Figure 2C** using cellular deconvolution analysis. Rows represent genes and columns represent the cluster identified in the t-SNE plot (bottom). Not all clusters have highly discriminatory unique markers. Only clusters having highly discriminatory markers are shown. For a cluster with more than 50 unique markers, only the top 50 markers are shown. The complete list of markers and the reference basis matrix used to perform deconvolution analysis are provided in supplementary table S4.

### **SUPPLEMENTAL TABLES** (each as separate excel file) with detailed notes

**Table S1:** Complete list of differentially expressed genes (Deseq2 output)

- **Worksheet 1 [DEGs]:** Complete list of genes with their differential expression profile. The differential expression analysis was performed using Deseq2 package in R.
- **Worksheet 2 [DEG_Bins]:** Differentially expressed genes with adjusted p-value < 0.05. The genes are ordered based on different log fold change bins used in the analysis and are colored as per Figure 1B.

Details for both worksheets:

**Gene_ID:** Ensembl ID for each gene.

**Gene_Names:** Gene symbol for each gene Ensembl ID.

**baseMean:** Average of normalized gene count.

**Log2FoldChange:** Log_2_ fold change. The effect size estimate. Positive value: upregulated in epilepsy with respect to control. Negative value: downregulated in epilepsy with respect to control.

**lfcSE:** The standard error estimate of the log_2_ fold estimate.

**Stat:** Wald statistics for Epilepsy vs Control comparison.

**pvalue:** Wald test p-value for Epilepsy vs Control comparison

**padj:** Benjamini-Hochberg adjusted p-value.

**Table S2:** Pathways associated with the differentially expressed gene bins (Corresponds to Figure 1C)

- **Worksheet 1 [-log10(FDR)]:** -log_10_(FDR < 0.05) of each enriched gene ontology pathway in different bins of log fold change is shown. Blue: downregulated; orange: upregulated.

Details Worksheet 1 [-log10(FDR)]:

**Pathways:** Gene ontology pathways with their respective IDs.

**Column C to K:** Different bins of log2 fold change corresponding to Figure 1B.

**Themes:** Different biological themes to which each pathway belongs.

#### **Table S3:** Predictive gene and associated pathways

- **Worksheet 1 [PredictiveGenes]:** List of genes predictive of the epilepsy phenotype using the sparse classifier. Genes are organized by down- and up-regulated.
- **Worksheet 2 [Pathways_Up]:** Pathway analysis of the up-regulated predictive genes using Gene Ontology.
- **Worksheet 3 [Pathways_Down]:** Pathway analysis of the down-regulated predictive genes using Gene Ontology.

Details for Worksheet 1 [PredictiveGenes]:

Subset of Deseq2 output (from Table S1, Worksheet 1 [DEGs]) for 119 predictive genes. Details of each column same as Table S1.

Details for Worksheet 2 and 3 (output from gene ontology database):

**Source:** different gene-ontology category to which the down- and up-regulated pathways belong. CC: Cellular Component, BP: Biological Process and MF: Molecular Function.

**Pathways:** Gene ontology pathways with their respective IDs.

**Mus musculus – REFLIST (22265):** Number of genes from the mouse reference gene list that map to a given pathway.

**Upload_1 (62)/ (59):** Number of genes in the predictive gene list (up: 62 and down: 59) that map to a given pathway.

**Expected:** Based on the *M. musculus* REFLIST, the number of genes expected in the predictive gene list (down- and up-regulated) for a given pathway.

**Over/Under:** + sign represents the over representation of a pathway in the predictive gene list (down- and up-regulated pathways are listed in separate worksheets).

**Fold Enrichment:** Number in column “Upload_1 (up: 62)/(down:59)” divided number in column “Expected”, i.e. fold enrichment of genes observed in predictive gene list (up and down separately) over the expected value.

**P-value:** Raw p-value as determined by Fisher’s exact test.

**FDR:** Benjamini-Hochberg adjusted p-value.

#### **Table S4:** Results for single cell RNAseq and deconvolution analysis

- **Worksheet 1 [Seurat_Output]:** List of genes in each respective cell cluster, as indicated in the column “Cluster.”
- **Worksheet 2 [Discrete_Markers]:** List of genes used to discriminately identify each cell-cluster (corresponds to Figure S2).
- **Worksheet 3 [S_Matrix]:** The gene expression signature matrix.
- **Worksheet 4 [Cibersort_Output]:** Proportional expression value of discrete markers in each individual biological sample (rows) across each cell cluster (columns). This data was used to generate Figure 2C.

Details for Worksheet 1 [Seurat_Output]:

**gene:** Gene symbol for each gene.

**p_val:** unadjusted p-value.

**avg_logFC:** Fold change of the average expression between the two groups (cluster in which the gene is differentially expressed as compared to remaining clusters). Positive values indicate that the feature is more highly expressed in the cluster assigned to it.

**pct.1:** Percent of cells where the gene is detected in the assigned cluster.

**pct.2:** Percent of cells where the gene is detected in the remaining cluster.

**p_val_adj:** Adjusted p-value, based on Bonferroni correction using all genes in the dataset.

**cluster:** Cluster in which the gene is significantly upregulated.

Details for Worksheet 2 [Discrete_Markers]: List of genes which are expressed 3 times higher in a given cluster as compared to other clusters. As compared to being differentially expressed or enriched, these genes are highly exclusive to a given cluster. These genes are used to generate the gene expression signature matrix **(Worksheet 3 [S_Matrix])** and perform the deconvolution analysis **(Figure 2C)** and make the heatmap in **Figure S2.**

Details for Worksheet 3 [S_Matrix]: Group of genes (rows) whose expression values collectively define a unique gene-expression signature(Newman et al., 2015) for each cluster (columns) and was used to deconvolve the bulk RNAseq. The matrix is presented as a heatmap in **Figure S2.**

Details for Worksheet 4 [Cibersort_Output]: The column represents cell-type clusters from S_Matrix (Worksheet 3) and rows represent deconvolution results for each sample. All results are reported as relative fractions normalized to 1 across all cell-type clusters. This output was used to generate **Figure 2B.**

Metrics for each sample (Cibersort_Output):

**P-value:** Statistical significance of the deconvolution results across all the cell-type clusters.

**Correlation:** Pearson's correlation coefficient (R), generated from comparing the original bulk RNAseq expression profile with the estimated expression profile. The estimated expression profile is calculated using imputed cell fractions and corresponding expression profiles from the signature genes file. Note that the correlation is restricted to discrete markers.

**RMSE:** Root mean squared error between the original bulk RNAseq expression profile and the estimated expression profile, restricted to restricted to discrete markers.

**Table S5:** Genes and pathways influenced by altered proportion of pyramidal neurons, interneurons, and non-neuronal cells.

- **Worksheet 1 [PYC-influenced]:** Complete list of differentially expressed genes influenced by all the pyramidal neurons.
- **Worksheet 2 [Int-influenced]:** Complete list of differentially expressed genes influenced by all the interneurons.
- **Worksheet 3 [NN-influenced]:** Complete list of differentially expressed genes influenced by all the neuronal cells.
- **Worksheet 4 [Influenced Pathways]:** Up- and down-regulated GO pathways associated with genes influenced by different cell-types (PYC, Int, and NN from worksheet 1, 2 and 3 respectively).
- **Worksheet 5 [Figure 3B]:** Up and down regulated GO pathways per cell type (from Worksheet 4) filtered by themes presented in Figure 1C. (corresponds to Figure 3B).

Details for Worksheet 1 to 3:

**Gene_ID:** Ensembl ID for each gene.

**Gene_Names:** Gene symbol for each gene Ensembl ID.

**baseMean:** Average of normalized gene count.

**LogFC:** Log_2_ fold change after adjusting for altered proportion of pyramidal neurons, interneurons, and non-neuronal cells. The effect size estimate. Positive value: upregulated in epilepsy with respect to control. Negative value: downregulated in epilepsy with respect to control.

**basePadj:** Benjamini-Hochberg adjusted p-value for epilepsy vs control comparison (i.e. without adjusting for altered proportion of pyramidal neurons, interneurons, and non-neuronal cells).

**adjustedpadj:** Benjamini-Hochberg adjusted p-value for epilepsy vs control comparison after adjusting the altered proportion of pyramidal neurons, interneurons, and non-neuronal cells.

**Regulated**: A gene with further reduced p-value (after adjusting for altered proportion of pyramidal neurons, interneurons, and non-neuronal cells) is considered to be influenced by alerted cell type proportion. TRUE= adjustedpadj < basePadj. FALSE= adjustedpadj > basePadj.

Details for Worksheet 4 and 5: All results are reported as -log10(FDR corrected P-value). This output was used to generate Figure 3B.

**Pathways:** Gene ontology pathways with their respective IDs.

**Column C to H:** Up and down regulated pathways for pyramidal neurons, Interneurons and non-neuronal cells.

**Themes (for Worksheet 5 only):** Different themes to which each pathway belongs. Themes taken from Figure 1C.

#### **Table S6:** WGCNA modules used to construct the Bayesian network and their gene ontology characterization

- **Worksheet 1 [WGCNA_Modules]:** Complete list of genes in each module for the Bayesian network.
- **Worksheet 2 [Module_Characterization]:** Characterization of each module by GO with BP, MF, and CC terms.
- **Worksheet 3 [Eigengene]:** Matrix of inferred eigengenes (Pigengene output).

Details of Worksheet 1 [WGCNA_Modules]:

**GeneNames:** Gene symbol for gene used to create the Bayesian network.

**Module:** WGNCA module to which the gene belongs to.

Details of Worksheet 2[Module_Characterization]:

**Module:** WGCNA module (only those used to create the Bayesian network).

**Source:** different gene-ontology category to which the down- and up-regulated pathways belong. CC: Cellular Component, BP: Biological Process and MF: Molecular Function.

**Pathways:** Gene ontology pathways with their respective IDs.

**Mus musculus – REFLIST (22265):** Number of genes from the mouse reference gene list that map to a given pathway.

**Query:** Number of genes in WGCNA module that map to a given pathway.

**Expected:** Based on the REFLIST for mouse, the number of genes expected in the WGCNA module for a given pathway

**Over/Under:** + sign represents the over representation of the pathway in WGCNA module.

**Fold Enrichment:** Numbers in column “Query” divided numbers in column “Expected”, i.e. fold enrichment of genes observed in a WGCNA module over the expected value.

**P-value:** Raw p-value as determined by Fisher’s exact test.

**FDR:** Benjamini-Hochberg adjusted p-value

Details of Worksheet 2[Eigengene]: Each row corresponds to a sample and each column represents an eigengene. This Matrix was used to fit a decision tree classifier.

#### **Table S7:** The Connectivity Map probe list

Details:

The differentially expressed up and down regulated genes lists belonging to different WGCNA modules connected to source and sink nodes. The directions (up and down) of the list were inferred using the effect size estimate (Log_2_ fold change) from Deseq2 output **(Table S1)**. Top Differentially expressed genes and directionally segregated predictive gene list **(Table S3, Worksheet 1[PredictiveGenes]** were also used as probe.

**Table S8:** Comprehensive list of pro-disease and therapeutic drugs from the connectivity map enriched in up- and down-stream nodes (WGCNA modules), predictive gene list, and differentially expressed genes.

- **Worksheet 1 [MOA_Prodisease]:** Complete list of drugs that are concordant with the disease signature in source and sink nodes **(Figure 4 A and B)**.
- **Worksheet 2 [MOA_Therapeutic]:** Complete list of drugs that are discordant with the disease signature in source and sink nodes **(Figure 4 A and B)**.
- **Worksheet 3 [Targets]:** The unique and common targets of prodisease and therapeutic drugs are listed.
- **Worksheet 4 [Definitions]:** Details of all the unique and common drug targets is provided.

Details for Worksheet 1 and 2:

**Molecules:** Name of the molecules whose gene-signatures are either concordant (prodisease, Worksheet 1) or discordant (therapeutic, Worksheet 1) with the source and sink node associated gene list (Table S7).

**Column C to J:** Source and sink associated nodes (M04, M05, M07, M10, M12 and M47), PG: Predictive gene list, and TopDE: Top differentially expressed genes.

**Description:** Mode of action associated with the molecule.

**Target:** Protein that the drug molecule targets.

-log_10_(q-value) of the hypergeometric overlap between the drug signature (from cmap) and source and sink node associated gene signatures (Table S7) is shown. Data for selected molecules are plotted in **Figure 5**.

Details for Worksheet 3 [Targets]: List of pro disease and therapeutic targets. Overlapping list elements are listed as common target whereas the non-overlapping ones are listed as unique.

Details for Worksheet 3 [Definitions]:

**Direction:** Targets classified as Common, Unique_Prodisease or Unique_Therapeutic (form worksheet 3).

**Target:** Targets form worksheet 3.

**Type:** Biological classification of the target molecule.

**Definition:** Details regarding the target molecule gathered from various internet resources.

**REFERENCE:**

Newman AM, Liu CL, Green MR, Gentles AJ, Feng W, Xu Y, Hoang CD, Diehn M, Alizadeh AA. 2015. Robust enumeration of cell subsets from tissue expression profiles. *Nat Methods* **12**:453–457. doi:10.1038/nmeth.3337

Saunders A, Macosko EZ, Wysoker A, Goldman M, Krienen FM, de Rivera H, Bien E, Baum M, Bortolin L, Wang S, Goeva A, Nemesh J, Kamitaki N, Brumbaugh S, Kulp D, McCarroll SA. 2018. Molecular Diversity and Specializations among the Cells of the Adult Mouse Brain. *Cell*. doi:10.1016/j.cell.2018.07.028
